# Metabolic complexity drives divergence in microbial communities

**DOI:** 10.1101/2023.08.03.551516

**Authors:** Michael Silverstein, Jennifer M. Bhatnagar, Daniel Segrè

## Abstract

Microbial communities are shaped by the metabolites available in their environment, but the principles that govern whether different communities will converge or diverge in any given condition remain unknown, posing fundamental questions about the feasibility of microbiome engineering. To this end, we studied the longitudinal assembly dynamics of a set of natural microbial communities grown in laboratory conditions of increasing metabolic complexity. We found that different microbial communities tend to become similar to each other when grown in metabolically simple conditions, but diverge in composition as the metabolic complexity of the environment increases, a phenomenon we refer to as the divergence-complexity effect. A comparative analysis of these communities revealed that this divergence is driven by community diversity and by the diverse assortment of specialist taxa capable of degrading complex metabolites. An ecological model of community dynamics indicates that the hierarchical structure of metabolism itself, where complex molecules are enzymatically degraded into progressively smaller ones, is necessary and sufficient to recapitulate all of our experimental observations. In addition to pointing to a fundamental principle of community assembly, the divergence-complexity effect has important implications for microbiome engineering applications, as it can provide insight into which environments support multiple community states, enabling the search for desired ecosystem functions.

## Introduction

Understanding how complex microbial communities assemble is important for addressing open challenges in microbial ecology with applications that range from medicine^1,2^ to climate change mitigation^3–5^. Studies in natural^6,7^ and laboratory^8–11^ settings have investigated the reproducibility of assembly dynamics across a range of environmental conditions leading to seemingly contradictory results. Under certain conditions, microbial community assembly appears to be highly deterministic, as different communities are driven by strong environmental selection towards a specific steady state independent of their initial composition^8^. Under other conditions, however, environmental selection is weaker, resulting in highly variable assembly of communities with more dependence on their initial composition^9^. Uncovering what properties govern this variability in microbial community assembly constitutes one of the fundamental questions of microbial ecology^12^ and is crucial for successful microbiome engineering, which aims to steer communities towards a desired structure in a given environment^13^.

Here, we combined experimental measurements and computational modeling to investigate the interplay of initial composition and environmental selection in determining community assembly and its variability. We followed the dynamic assembly of diverse microbial communities inoculated from different soil samples grown on carbon sources of increasing metabolic complexity. By tracking how closely these communities resembled each other over time, we found that the effect of environmental selection on communities depended on the metabolic complexity of the environment itself. Specifically, different microbial communities diverged in their taxonomic composition across a gradient of increasingly complex metabolic conditions, suggesting that the forces dominating microbial community assembly shift from strong to weak environmental selection in increasingly complex conditions. By constructing a consumer resource model that recapitulates this effect, we additionally learned that this divergence-complexity relationship depends on a hierarchical structure of metabolite transformations (e.g. polysaccharides to oligosaccharides to monosaccharides), but does not depend on the distribution of these metabolic functions across taxa. Our results point to an ecosystem organization principle that can help reconcile seemingly incompatible observations of divergence in different conditions and provide guidelines for which environments may be more susceptible to microbiome engineering projects.

### The divergence-complexity effect hypothesis

To assess the strength of environmental selection on community assembly, one would ideally compare how the trajectories of multiple distinct microbial communities diverge in taxonomic composition across a set of conditions. A key question we ask is whether distinct communities assembled in the same condition tend to become taxonomically similar and how the degree of similarity depends on the metabolic complexity of the environment. For example, we can imagine how different microbial communities that initially vary in taxonomic composition (**Fig. 1a**) may converge in composition over time when grown in one environment (strong environmental selection, **Fig. 1b**) while those same communities may diverge in another environment, arriving at alternative stable states (weak environmental selection, **Fig. 1c**). To quantify the degree to which different communities diverge taxonomically from each other when grown in a given condition, we calculate the difference (beta diversity) in their compositions as they develop towards a steady state (Methods; **Fig. 1d**).

**Fig. 1.**
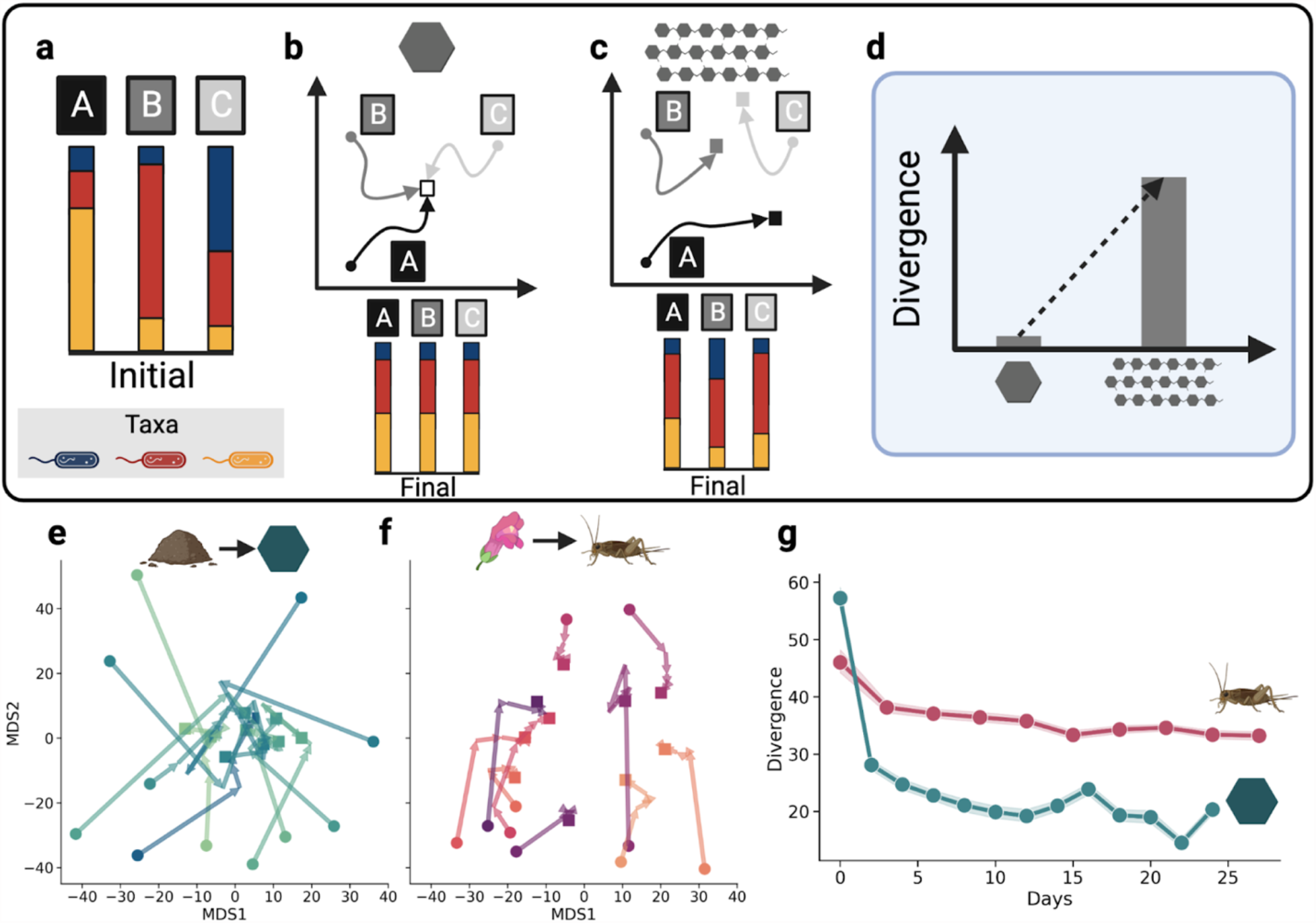
Microbial communities may diverge in environments with increasing metabolic complexity. **a**-**d**, Hypothesis of microbial community divergence in theoretical simple (**b**) and complex (**c**) metabolic conditions. Microbial communities A, B, and C are initially composed of different compositions of the same three microbial species (**a**; blue, red, and yellow). Over time, communities grown on a simple substrate (**b**) converge, while these same communities grown on a complex substrate (**c**) diverge. **d**, Quantification of divergence at the final time point for hypothetical scenarios in **a** and **b. e-g**, Divergence observed in two independent experimental studies where microbial communities were sourced from soils or leaves and grown on glucose (**e**; a relatively simple metabolic environment from Goldford et al.) and communities were sourced from pitcher plants and grown on acidified cricket media (**f**; a more complex metabolic environment from Bittleston et al.). Each colored line in **e** and **f** represents the trajectory of a community’s composition over time in separately computed multidimensional scaling (MDS) projections. **g**, The divergence for each metabolic environment, calculated as the pairwise distances between all communities within a given condition at each time point. Each point is the mean pairwise distance within condition at each time point and shading represents the 95% confidence interval over all pairwise distances within each environment at each timepoint.

We initially identified existing data that could indicate whether and how community divergence would indeed depend on environmental conditions. We re-analyzed two independent studies that both explored how a collection of diverse microbial communities assembled over time, but did so under very different conditions. When one study, Goldford et al.^8^, cultured communities in (simple) glucose media, communities converged (**Fig. 1e**). By contrast, when Bittleston et al.^9^ cultured communities in (complex) acidified cricket media, they diverged (**Fig. 1f**). In both cases, the initial communities differed substantially from each other and then immediately became more similar; however, communities enriched on glucose ultimately converged significantly more, despite starting with greater variation in initial community composition (**Fig. 1g**). Based on the striking discrepancy in the degree of divergence across these two studies we formulated the hypothesis that divergence increases with the metabolic complexity of the provided resources (**Fig. 1d**), a relationship that we will refer to as the divergence-complexity effect.

### Community divergence increases with metabolic complexity

In order to directly test the divergence-complexity effect, we designed an experiment to quantify the divergence of microbial communities grown in conditions of increasing metabolic complexity (Methods; **Fig. 2a**). To assess divergence, we sourced six microbial communities from forest soils that are generally diverse and distinct from each other^14^, even over small (centimeter) spatial scales^7^. Each microbial community was grown in nine different minimal media, each supplemented with equimolar concentrations of at least one carbon source commonly found in soils^15^: (1) citrate, (2) glucose, (3) cellobiose, (4) cellulose, (5) lignin, (6) citrate + glucose, (7) citrate + glucose + cellobiose, (8) citrate + glucose + cellobiose + cellulose, or (9) citrate + glucose + cellobiose + cellulose + lignin. In testing the divergence-complexity effect, we consider metabolic complexity to increase from citrate to lignin (in line with the number of metabolic byproducts expected from each metabolite^16^). We included single- and mixed-metabolite conditions in order to test the divergence-complexity effect with increasing complexity of each metabolite (single), as well as increasing resource diversity (mixed). Each microcosm, containing one source community growing in one condition, was serially passaged ten times, in intervals of three days. 16S rRNA sequencing was performed and amplicon sequence variant (ASV) counts were generated for the initial soil inocula and microcosm communities at days 3, 6, 9, 12, and 33.

**Fig. 2.**
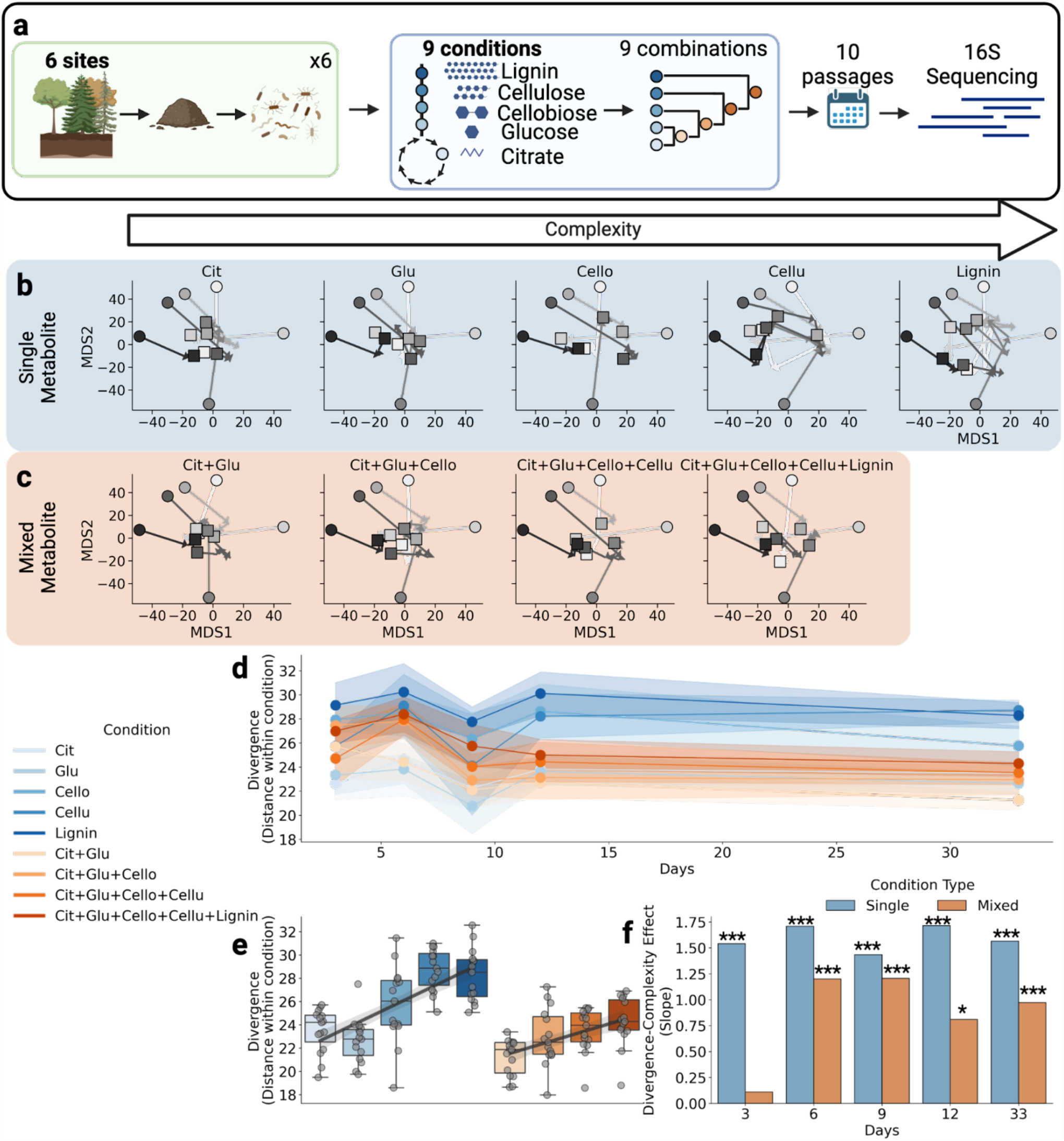
Divergence of microbial communities increases in environments of increasing metabolic complexity. **a**, Study design: microbial communities were extracted from six forest soils and were then grown in nine conditions (citrate, glucose, cellobiose, cellulose, lignin, citrate + glucose, citrate + glucose + cellobiose, citrate + glucose + cellobiose + cellulose, and citrate + glucose + cellobiose + cellulose + lignin). Communities were passaged ten times once every three days and sequenced on days 0, 3, 6, 9, 12, and 33. **b-c**, MDS projections of community trajectories over time in each single-metabolite condition (**b**) and mixed-metabolite condition (**c**). MDS was calculated on all samples together for ease of visually comparing trajectories between conditions. **d**, Divergence of communities within each condition over time from day 3 onwards. Initial communities are a distance of 58.1+/-3.5 (not shown for clarity). Single metabolite conditions are in blue, mixed conditions are in orange, and colors darken with complexity. Points on each line represent the mean divergence and the shaded region represents the 95% confidence interval for pairwise distances between all six communities within each condition. **e**, Distribution of divergence for the final time point where divergence increases with metabolic complexity for single and mixed-metabolite conditions (same colors as **d**). **f**, Metabolic complexity effect by condition type (slopes shown in **e**) for all time points. P-values computed on significance of effect (slope) > 0 (*: p<.05, ***: p<1e-3).

Supporting our hypothesis of the divergence-complexity effect, we observed that divergence increased with metabolic complexity (**Fig. 2b-f**). In accordance with previous studies (**Fig. 1e-f**), our source communities initially differed from each other and then immediately converged and stabilized once introduced to laboratory conditions (**Supp. Fig. 1, Fig. 2d**). Once stabilized, we observed the divergence-complexity effect on single- and mixed-metabolite conditions, separately (**Fig. 2e**). Within single-metabolite conditions, the communities converged strongly on simple metabolites, while they diverged to increasingly distinct states on the more complex metabolites (**Fig. 2b, 2d-f**). Similarly, community divergence increased from the least (citrate + glucose) to the most diverse (all metabolites) mixed-metabolite conditions (**Fig. 2c-f**). Interestingly, the effect is stronger in single-metabolite conditions than in mixed-metabolite conditions (**Fig. 2e**), suggesting that assembly dynamics are sensitive to the order in which different metabolites become available through trophic interactions^17^. These trends are detectable at each sampled time point (**Fig. 2f**) and when we re-computed divergence at the Family taxonomic level (**Supp. Fig. 3-4**). Because bacteria often differ in metabolic function at the Family level^18^, this latter result suggests that our communities, which assemble to distinct taxonomic compositions, may also be engaging in distinct metabolic activities.

The degree of divergence in complex conditions appears to be particularly sensitive to differences in initial community composition. Communities sourced from different locations, but that were initially similar to each other, did not necessarily converge to similar final states (**Fig. 2b-c**), suggesting that they may be traversing a rugged structure-function landscape in complex conditions, where slight differences in initial composition can lead to distinct final compositions^19^. Conversely, replicate microcosms assembled from the same source did cluster together (**Supp. Fig. 2**), suggesting that while assembly dynamics are indeed complex (initially similar communities can diverge), they are reproducible and not diverging merely due to stochasticity. These trajectories show that complex environments can support a greater number of discrete alternative stable states than simple environments.

### Community diversity dynamically correlates with divergence and implicates the role of specialists

In order to gain a deeper understanding of the divergence-complexity effect, we investigated how alpha diversity within each individual community correlates with divergence across communities. In particular, two separate principles could jointly give rise to the divergence-complexity effect. The first principle, “metabolic complexity begets diversity”, where community diversity increases with increasing metabolic complexity, has been experimentally documented in both natural and synthetic communities^11,16^. A proposed second principle, “diversity begets divergence”, could result from the expectation that more diverse communities have more variation in the abundance of each microbe, leading to higher divergence across communities. If metabolic complexity yields diversity and diversity yields divergence, we would expect higher divergence in increasingly complex conditions, leading to the divergence-complexity effect.

Consistent with these expectations, we observed a strong linear relationship between diversity and divergence, which strengthened over time, indicating that specific changes in community assembly drive the rise of divergence. The slope of the diversity-divergence relationship increased over time (**Fig. 3a-b**), despite the fact that diversity itself, on average, decreased (**Fig. 3a**). In other words, over time, the same degree of divergence is maintained by communities with reduced diversity. For divergence to remain relatively stable while diversity decreases (**Fig. 2d**), taxa endemic (i.e. specific) to each community must persist while a set of species shared across communities universally go extinct within each condition. One possible explanation is that these persistent taxa are metabolic specialists, which produce enzymes that target specific biochemical bonds^20^. We hypothesize that functionally redundant^21^ specialists that differ between communities and target complex metabolites are less evenly distributed across communities than taxa that specialize on simpler metabolites, driving the divergence-complexity effect.

**Fig. 3.**
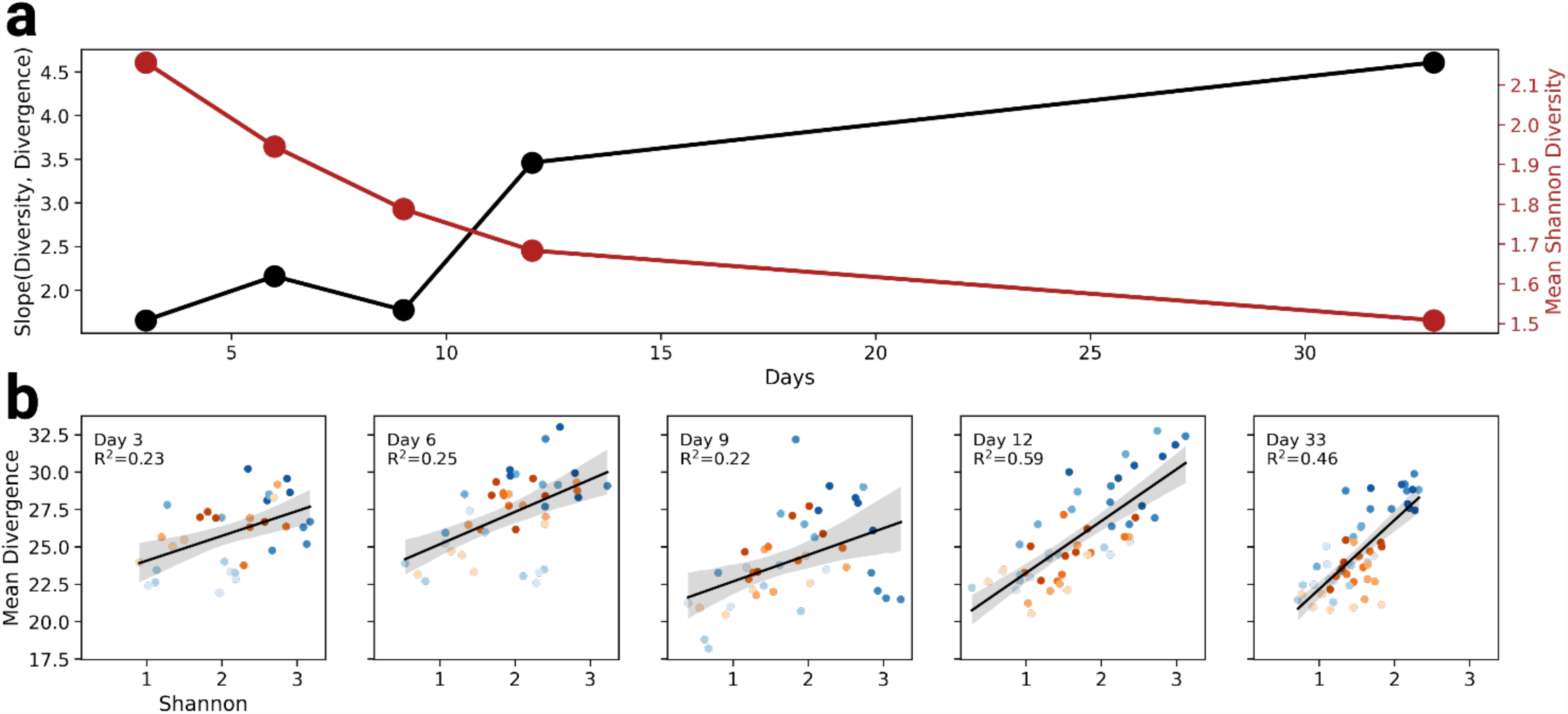
Divergence dynamically correlates with diversity. **a**, The slope of the relationship between community alpha diversity and divergence (red) and the mean community alpha diversity (gray) over time. Shaded areas around each regression line represents the 95% confidence interval. **b**, The data underlying the relationship in **a** over time. Each point is the diversity of a community in a condition (x-axis) and the divergence of that community from all others within a condition. While diversity is expected to increase with metabolic complexity, it is not clear if different communities will increase in diversity in the same ways.

### Specialists are more endemic in complex conditions

To investigate the role of specialists in the divergence-complexity effect, we explored the distribution of taxa across experimental conditions and source communities. If specialists drive the diversity-complexity effect, we would expect to see that specialists are increasingly endemic, or unevenly distributed across source communities, in more complex conditions.

To quantify the degree of specialization, we computed a condition-specificity metric for each taxon (ASV) in each condition, and then assessed whether specialization and endemism depended on metabolic complexity. We defined condition-specificity for each taxon and condition as the fraction of source communities in which that taxon was found at the final sampling time point on that given condition. In particular, if a taxon only occurs in one condition, its condition-specificity is 1 and will be referred to as a “specialist”. In accordance with our expectations, we observed that more complex conditions (particularly single-metabolite ones) had greater condition-specificity (**Fig. 4a**) and more specialists (**Fig. 4b**).

**Fig. 4.**
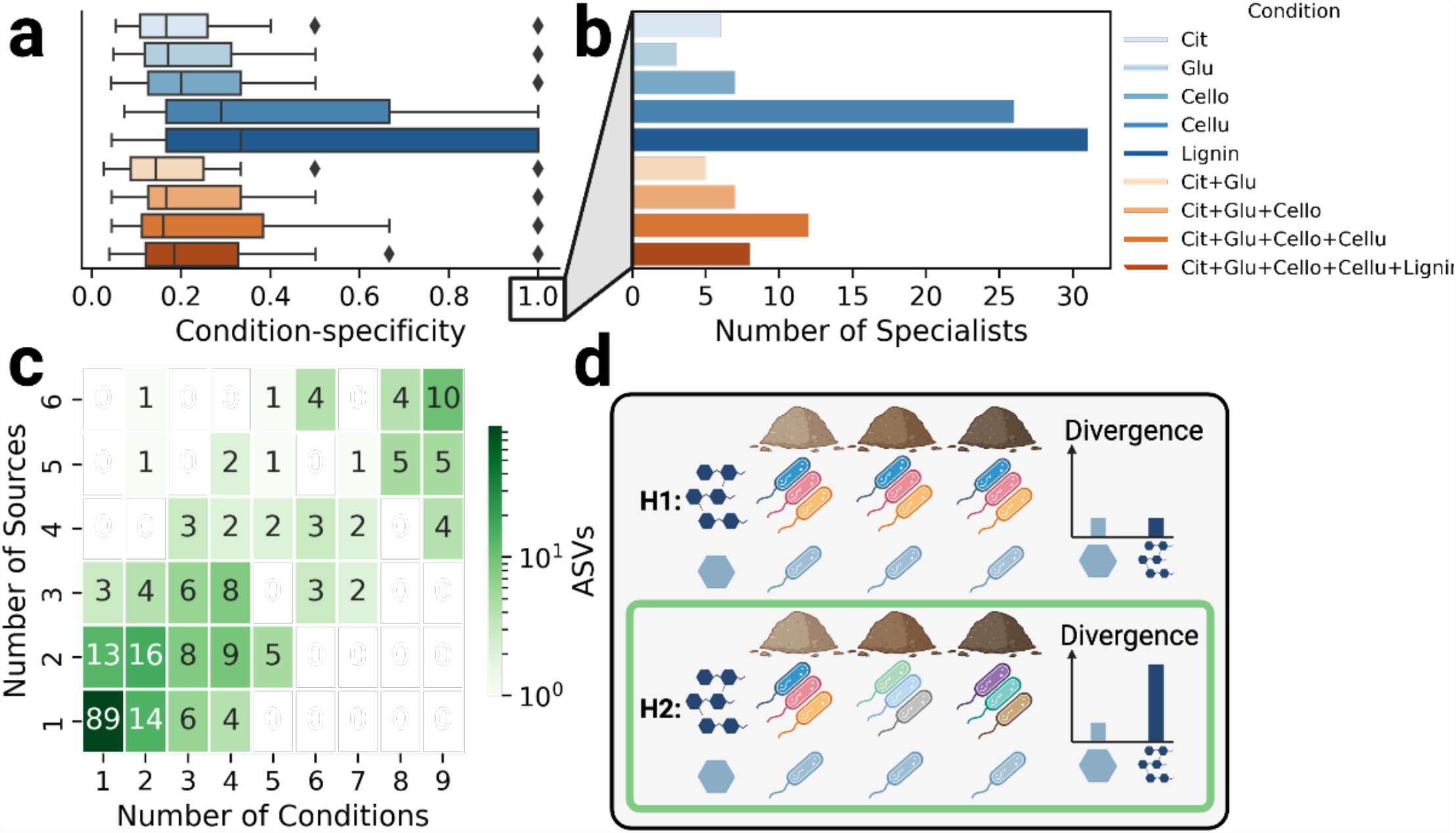
Endemic taxa are enriched and unevenly distributed in complex conditions. **a**, The distribution of condition-specificity per condition for day 33. Condition-specificity is calculated as the fraction of occurrences of a taxon that is attributed to a particular condition, such that a specificity of 1 means that taxon occurs in only one condition (a specialist). **b**, The number of specialists per condition. **c**, Taxon occurrence by number of conditions and number of source communities. ASVs found in fewer conditions are less evenly distributed across source communities (found in fewer source communities) and taxa found in more conditions are more evenly distributed across source communities. **d**, Two hypotheses for single metabolite conditions following from **a** and **b**, where H2 is supported and H1 is not. H1: more complex conditions are enriched for specialists and when those taxa are evenly distributed across source communities, it results in similar divergence for complex and simple metabolic conditions. H2: when more complex conditions are enriched for specialists and these taxa are less evenly distributed across communities, more complex conditions result in greater divergence.

These results alone are encouraging, but are not sufficient for linking specialists to the divergence-complexity effect, which would additionally require specialists to differ between communities in the same condition. While we observed an enrichment of condition-specific taxa in complex conditions, it hypothetically could be the case that these same taxa were found across all source communities, in which case communities in complex conditions would not diverge more than those in simple conditions (H1 **Fig. 4d**). However, when we count the occurrence of each taxon in each condition and source community, we find that condition-specific taxa are also source community-specific (endemic; **Fig. 4c**). As a result, taxa that specialize on complex metabolites are less evenly distributed across communities than taxa that specialize on simpler metabolites, and are therefore heavily implicated in mediating the divergence-complexity effect (H2 **Fig. 4d**).

### Trophic resource transformations reproduce divergence with consumer resource models

To better understand what aspects of the organisms and their environment are necessary for the diversity-complexity effect, we performed a series of simulations with microbial consumer-resource models (CRMs; Methods)^22^. In particular, we wanted to corroborate our hypothesized mechanism of the diversity-complexity effect, that divergence correlates with metabolic complexity and emerges from endemism of specialist taxa. CRMs are dynamical ecological models where consumers are defined by the set of resources they prefer (consumer preferences) and resources are able to be transformed by consumers into other resources following consumption (resource transformations). Overlapping consumer preferences give rise to competition, while the exchange of secreted transformed products can generate cross-feeding interactions^22^. Taxonomic structure in CRMs can be represented by specifying “families” of consumers that have similar resource preferences (specialization; **Supp. Fig. 6c-d**). Metabolic structure can be represented by assuming that upon metabolization, resources of a given type transform into resources of another specific type in a hierarchical fashion^23^ (Supp. **Fig. 6a-b**). In natural ecosystems, consumer preferences and resource transformations are typically arranged in a trophic structure, where taxa specialize in the hierarchical consumption of environmentally available metabolites and cross-feed the resulting (simpler) byproducts to taxa at subsequently lower trophic levels^17^. In order to understand whether the divergence-complexity effect could emerge solely from these ecological forces (consumer preferences and resource transformations), we assumed physiological parameters such as rates of consumer growth, consumer maintenance, resource utilization, resource energy density, and leakage (the fraction of transformed resource that is secreted) to be uniform across all consumers and resources^22,23^.

To investigate the role of taxonomic and metabolic structure in community divergence, we closely mimicked our experimental design and measured divergence of simulated communities using four different CRM configurations that captured combinations of trophic structures of consumer preferences and resource transformations, as well as corresponding random controls, similar to those shown to be sufficient to reproduce a number of ecological properties^23^ (**Fig. 5, Supp. Fig 6**, Methods). Trophic consumer preferences were defined with “families” of specialists and generalists (**Supp. Fig 6c**). In line with our experimental observations (**Fig. 4**), we set the diversity of specialists to be proportional to the complexity of the resource type they prefer. Trophic resource transformations were defined such that complex resources successively transformed into simpler ones in a hierarchical fashion (**Supp. Fig. 6a**). Random controls of consumer preferences (**Supp. Fig 6d**) and resource transformations (**Supp. Fig 6b**) were also generated, where consumers preferred resources of any type and resources transformed into others of any type, respectively. Combinations of these four parameterizations led to the following four model configurations: trophic consumer preferences and resource transformations (fully structured; **Fig. 5a**), random preferences and trophic transformations (resource structured; **Fig. 5b**), trophic preferences and random transformations (consumer structured; **Fig. 5c**), and random preferences and transformations (fully random; **Fig. 5d**). All four model configurations were initialized with six source communities and seven conditions (four single and three mixed resource conditions) and growth dynamics were simulated until reaching a steady state.

**Fig. 5.**
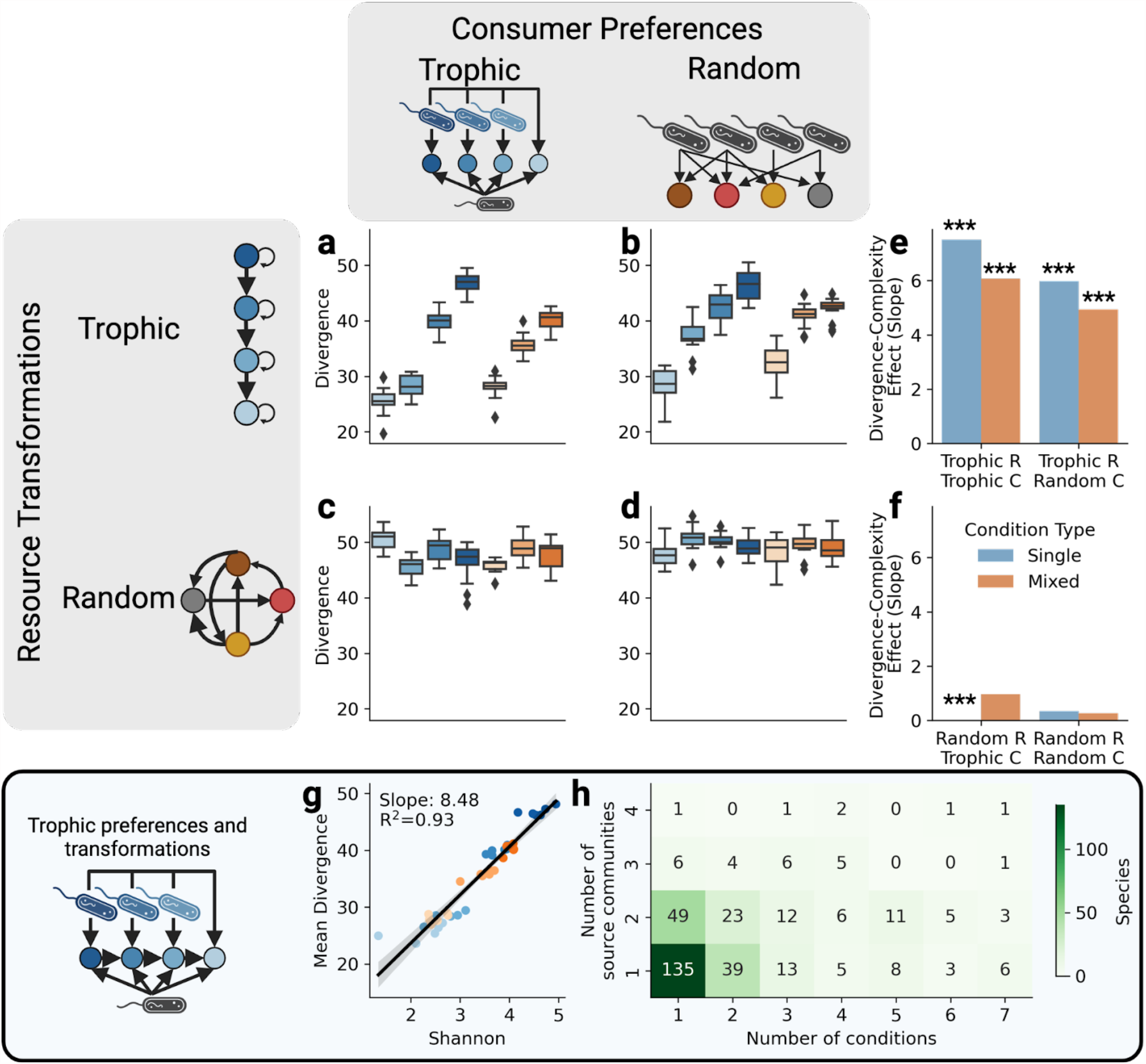
Trophic resource transformations reproduce divergence with consumer resource model simulations. The distribution of divergence for communities with simulated consumer resource models with and without trophic structure in resource transformations and consumer preferences. Mimicking our experiment, community growth was simulated in single metabolite (blue) and mixed metabolite (orange) conditions of increasing complexity (darker). **a-d**, Divergence for communities simulated with trophic resource transformations consumer preferences (fully structured; **a**), trophic resource transformations and random consumer preferences (resource structured; **b**), random resource transformations and trophic consumer preferences (consumer structured; **c**), and random resource transformations and consumer preferences (fully random; **d**). **e-f**, The effect of metabolic complexity on divergence for single and mixed metabolite conditions with trophic resource transformations (**e**) and random transformations (**f**; ***: p<1e-6). **g-h**, Using the fully structured configuration, the relationship between diversity and mean divergence (**g**) and the relationship between occupancy in conditions and number of source communities (**h**).

Surprisingly, our simulations showed that trophic structure of the resource transformations alone was necessary and sufficient to reproduce the divergence-complexity effect in both single- and mixed-resource conditions (**Fig. 5a-f**). Notably, even when consumer preferences were random, the divergence-complexity effect was still observed as long as resource transformations were structured (**Fig. 5a,b,e**). However whenever resource transformations were random, all communities diverged equally, irrespective of metabolic complexity, and thus there was no divergence-complexity effect (**Fig. 5c,d,f**). In addition to qualitatively reproducing the divergence-complexity effect, our model recovered, as emergent properties, further non-trivial trends detected in our experiment. For example, in model configurations with trophic resource transformations, the divergence-complexity effect is greater for single resource conditions than mixed ones (**Fig. 5e**), as observed experimentally (**Fig. 2f**). Additionally, the maximum divergence in single resource conditions exceeds that of mixed conditions (**Fig. 5a-b; Fig. 2e**). These model configurations also reproduced our downstream analyses, such as the correlation between divergence and diversity (**Fig. 5g; Fig. 3b**) and the tendency for specialists to be endemic (**Fig. 5h; Fig. 4d**). The reproduction of these patterns with physiologically neutral consumers (uniform physiological parameters) and resources implicates the trophic metabolic structure in resource transformations as the driving mechanism of the divergence-complexity effect.

## Discussion

Compelled by recent experiments which found that microbial community diversity increases with metabolic complexity^11,16^, we sought to reconcile contradictory interpretations of whether microbial communities tend to converge^8^ or diverge^9^ in the same conditions. By jointly revisiting these two propositions, we uncovered a new, reproducible, and quantitative ecological principle, the divergence-complexity effect, which has important consequences for ecological theory and microbiome engineering. While previous work explored community assembly by modulating the complexity of metabolic conditions^11,16,24^ or the variability of source communities^8–10^, the divergence-complexity effect could be observed only by systematically varying both, i.e. analyzing multiple source communities under increasingly complex conditions. We found that divergence correlates strongly with diversity, which is driven by an enrichment of specialists in complex conditions. We concluded our analysis by reproducing these results using consumer resource model simulations, which provide insights into the potential ecological mechanisms of the divergence-complexity effect.

While our experimental results are robust and reproducible, they necessarily rely on specific design constraints. Experimental choices that could be revisited in future studies include the passaging time, chosen here to be three days, as used in other microbial community assembly studies with complex metabolites^9^; the selection of metabolites, which constitute a representative, but oversimplified version of the metabolic complexity of soil environments; and the focus on taxonomic divergence (through 16S amplicon sequencing) rather than functional divergence, which would require a comprehensive profiling of microbial functions with metagenomic or metatranscriptomic sequencing. We designed our simulations to represent the ecological structure of microbial communities and organization of metabolites as accurately as possible; however, our consumer resource models lacked the encoding of certain granular processes such as diffusion, transcriptional regulation, and anti-microbial defense. Inclusion of these processes with other ecological models^25^ could help to reveal further mechanistic insights into the diversity-complexity effect.

The most surprising result from our simulations was how structured resource transformations (where complex metabolites are progressively degraded into simpler ones), but not consumer preferences, were required for reproducing the divergence-complexity effect (**Fig. 5**). This result disappears completely when the resource transformations are uniformly random. A possible interpretation of this result is that microbial community assembly and dynamics are strongly dependent on the actual structured architecture of metabolism, which differs substantially from a network of random transformations. We cannot rule out the possibility that adding more parameters, and increasing the realism of simulations may affect our results. For example, we could parameterize our models to incorporate the trade-offs that are known to exist between enzyme production and growth rate in nutrient limited conditions^26^. However, since our current model captures so many of our observations, including the similarities and subtle differences between the single- and mixed-metabolite conditions, it lends confidence to the dominant role that the architecture of metabolism plays in community structure, corroborating previous reports^16^.

Importantly, the divergence-complexity effect has direct implications for the engineering of microbial communities towards any target, suggesting that metabolically complex environments may be more susceptible to microbiome engineering than simple ones. Potential targets for microbiome engineering include correcting the dysbiosis in the human gut^2^ and increasing the carbon stabilization capacity of soils^5^, among many other microbially-regulated traits. The consequences of the diversity-complexity effect are encouraging for efforts along these lines, since complex environments may be more likely to support an alternative community that is equally stable as the original one, but with potentially increased expression of a trait of interest. Culturing techniques such as directed evolution, where a set of microbial communities undergoes iterative rounds of perturbation and artificial selection in order to assemble high-performing communities^27^, offer an ideal strategy for exploring the different alternative states that a complex environment can support. Future research is required in order to understand how, in light of functional redundancy^28^, the divergence in taxonomic composition that we observe relates to divergence in functional composition, since modifying functional activity is commonly the goal of microbiome engineering efforts. Ultimately, we envisage that the awareness of the divergence-complexity effect may help microbial ecologists reframe the role of environmental selection in microbial community assembly and enable further research into the engineering of complex microbially-regulated environments.

## Methods

### Media preparation

Eleven different media were generated at equimolar (50mM) concentrations of carbon (C) in increasing levels of complexity. Stocks of citrate, glucose, cellobiose, cellulose, and lignin were generated at 1 mol C/L (1M C) and then sterilized. Citrate, glucose and cellobiose stocks were sterilized through 0.2um RapidFlow filters while cellulose and lignin stocks, whose particle sizes were too large for filters, were autoclaved. C source stocks were then mixed with M9 minimal media (5x M9 salts, 1M MgSO_4_, 1M CaCl_2_, 1x trace minerals)^29^ to form the following nine conditions, each made at a final concentration of 50mM C and with equal ratios of each C source: citrate, glucose, cellobiose, cellulose, lignin, citrate + glucose, citrate + glucose + cellobiose, citrate + glucose + cellobiose + cellulose, and citrate + glucose + cellobiose + cellulose + lignin. All media were stored in glass bottles, wrapped in foil, and stored at 4°C.

### Sample collection and microbial community extraction

On October 27, 2022 about half a pound of organic horizon soil (5-10cm deep) was collected from six sites at Harvard Forest in Petersham, MA. Two were pine dominated, two were hardwood dominated, and two were mixed. Samples were collected 15m from the forest edge and kept on ice until transported back to the laboratory the same day. Fresh soils were sieved through a 2mm mesh and then stored at 4°C. On November 21, 2022, 20g of each sieved soil was individually combined with 100mL of sodium pyrophosphate to separate cells from soils^30^ and was blended for three cycles of 10 seconds at ∼22,000 RPM (https://www.rosewill.com/rosewill-rhpb-18001-68-ounces-jar-size-1400w/p/9SIA072GJ93074) and then off for 10 seconds, and then 25mL of the resulting slurry was transferred to a centrifuge tube. The blender was washed between each sample by blending in 500mL of diluted bleach. Following the blending of all soils, each slurry was centrifuged for 10 minutes at 20,000xg, resuspended in 30mL of PBS, and rocked on an orbital shaker for 1 hour^16^ at 4°C. After rocking, samples were allowed to settle for 5 minutes and then passed through a 100um cell straining filter. Optical density (OD) measurements were performed at 600nm at a 1:20 dilution, 500uL of each sample was stored at -80°C in 20% glycerol, and the remaining volume from each sample used for inoculating experimental plates.

### Experimental culturing

Community extracts were added to 96-deep well plates in triplicate with all media combinations (3 replicates of each source community in each condition), generating a total of 162 microcosms (9 media combinations x 6 source communities x 3 replicates). Cycloheximide, an antifungal agent, was added to each well at 200ug/mL^31^ to reach a final OD of 0.1 and volume of 400uL per well. Plates were then stored in an incubator at 25°C under constant shaking at 200RPM. Communities were passaged every 72 hours into fresh media at a 1:20 dilution (without cycloheximide) for a final volume of 400uL, OD_600_ was measured, and the remaining volume was stored at -80°C for DNA extraction.

### DNA extraction and sequencing

DNA extraction for the six initial forest soil communities was performed using the PowerSoil DNA extraction (QIAGEN). After adding lysis buffer, samples underwent three cycles of freezing in liquid nitrogen, warming at 55°F in a water bath, and bead-beating for 1 minute (PowerLyzer, MoBio), then following the provided protocol for the remainder of the extraction. DNA was extracted from an additional 330 lab cultured samples on different days for each forest site and condition on days 3, 6, 9, 12, and 33, where available (**Supp. Table 1**). DNA extraction from lab cultured samples was performed using the PureLink Pro 96 Genomic DNA Kit (ThermoFisher) following the provided protocol except for extending all lysis incubation periods to two hours. DNA extracts were sent to Quintara Biosciences for library preparation and 16S amplicon sequencing using V4 primers 515F (GTGYCAGCMGCCGCGGTAA) and 806R (GGACTACNVGGGTWTCTAAT) on a single Illumina MiSeq run.

### Amplicon sequence processing

We received raw sequencing data for our study from Quintara Biosciences and downloaded raw sequencing data from Goldford et al.^8^ (SRP144982) and Bittleston et al.^9^ (SRP218147) from NCBI. All raw 16S sequencing data for each study was separately processed using BU16S (https://github.com/Boston-University-Microbiome-Initiative/BU16s), a QIIME2^32^ pipeline customized to run on Boston University’s Shared Computing Cluster. Briefly, BU16S first trims primers and filters out reads of less than 50 base pairs using cutadapt^33^, then obtains ASVs using dada2^34^, and finally classifies ASVs with 95% or greater sequence identity to the SILVA_132_99 database with VSEARCH^35^.

## Data analysis

All data analysis was performed in Python version 3.8.11. Pairwise distances between samples were computed with the Aitchison distance because it accounts for the compositional nature of sequencing data, unlike common distance metrics, such as Bray-Curtis, Jenson-Shannon Divergence, and Unifrac^36,37^. The Aitchison distance, A, between two compositions **x** and **y** is:

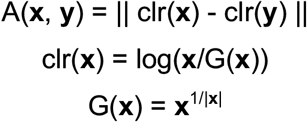

Where clr is the center-log ratio transform and G is the geometric mean. Divergence was computed by calculating the Aitchison distance between all pairs of samples within a condition at each timepoint. The divergence for samples at day 33, where we have up to three replicates for each community, is reported as the mean pairwise distance between all replicates. The mean divergence for each sample in a given condition is computed as the mean pairwise distance from each sample to all other samples in that condition.

Dimensionality reduction of pairwise distances was performed using multidimensional scaling (MDS) in scikit-learn^38^. MDS was computed separately for samples from Goldford et al. and Bittleston et al., while for our data MDS was computed jointly on all samples to allow for ease of comparability when viewing community trajectories in separate conditions.

Alpha diversity was computed by first rarefying (subsampling) all samples to 5028 reads and dropping twelve samples below this sequencing depth from subsequent alpha diversity analyses. The Shannon Diversity Index was calculated as -Σ**x**ln**x** and the ecological richness was calculated as Σ**x**>0 for each sample composition, **x**.

Condition specificity was calculated for each ASV by calculating the fraction of times each ASV was present in each condition. ASVs with a condition specificity of 1 were considered “specialists” since they were only found to occur in a single condition.

### Consumer resource models

We simulated the growth of 168 microcosms (6 communities x 7 conditions x 4 configurations) using microbial consumer resource models (CRM). With microbial CRMs, the dynamics of species and resources can be modeled with the following equations:

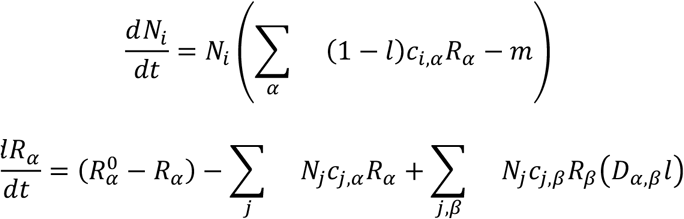

Where *N*_*i*_ is the abundance of species *i, R*_*α*_ is the concentration of resource *α*, 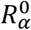 is the resource supply concentration, *l* is the leakage fraction i.e. how much each resource is “leaked” (how much of *α* is converted into *β*, were the rest is converted into biomass), *m* is the consumer maintenance cost, *c*_*i*,α_ is the consumer preference matrix, and *D*_*α, β*_ is the resource transformation matrix describing the rate that *β* turns into *α* following consumption^22^. We fixed the leakage (*l*=0.8), maintenance (*m*=1), and uptake rates (*c*_*i,α*_={0, 1}) for all consumers resulting in an “ecologically neutral”.

In order to study the impacts of trophic structure on divergence, we explored four different CRM configurations that varied in whether or not resource transformations or consumer preferences were trophically structured or random (**Supp. Fig. 6**). Resources were defined by establishing a resource pool of four resource types, T0, T1, T2, and T3, where each type consisted of 80, 60, 40, and 20 resources, respectively. Trophic resource transformations were parameterized by defining resources of one type to subsequently transform into resources of another type in a unidirectional fashion, with some self-renewal (**Supp. Fig. 6a**). For example, T0-type resources mostly transform into T1-type resources and some T0-type resources (**Supp. Fig. 6a**). Random resource transformations were defined by allowing each resource transform into any other resource with uniform probability (**Supp. Fig. 6b**). Transformation profiles for each resource in both configurations were sampled from Dirichlet distributions.

Consumers were defined by establishing a metacommunity of four “families”, F0, F1, F2, and G, where each type consisted of 500, 300, 100, and 100 consumers, respectively. Trophic consumer preferences were defined by allowing consumers of each family to utilize a total of 35 sampled resources from their associated type (i.e. F0 consumers could utilize T0 resources) and a common resource type. The skewed distribution of consumer family size was chosen to model our experimental results where the number of specialists correlated with metabolite complexity. Consumers belonging to the G (generalist) family could consume resources of any type (**Supp. Fig. 6c**). Random consumer preferences were defined by allowing each consumer to utilize 35 random resources from any type (**Supp. Fig. 6d**). Code for sampling trophic resource transformations and trophic consumer preferences can be found at: https://github.com/michaelsilverstein/ms_tools/blob/main/ms_tools/crm.py.

Initial conditions and source communities were defined to mimic our experimental design. Seven conditions were defined by sampling 20 resources from each resource type for single-metabolite conditions (T0, T1, T2, and T3) and from mixtures of resource types for mixed-metabolite conditions (T3+T2, T3+T2+T1, T3+T2+T1+T0; **Supp. Fig. 6e**). Six source communities were defined by sampling 200 consumers from the metacommunity (**Supp. Fig. 6f**).

The dynamics of each source community was then simulated in each condition using all four parameter configurations (trophic transformations and preferences, trophic transformations and random preferences, random transformations and trophic preferences, and random transformations and preferences) and the divergence was computed for each condition and configuration. Simulations of community assembly were performed by passing model parameters (D matrix, c matrix, and initial conditions) to the Community Simulator package^39^, which provides utility functions for constructing and solving the system of ordinary differential equations. In order to appropriately compare our simulation results, which simulates actual abundances of each consumer, to our experimental results, which reports relative abundance of each ASV, we rescaled the abundance of all communities to the same range to simulate the process of sequencing. Divergence was then calculated on the rescaled simulated community composition profiles in the same way as with our experimental data (using the Aitchison distance).

## Acknowledgements

We would like to thank Martina Dal Bello, Akshit Goyal, Matti Gralka, Zoey Werbin, Jing Zhang, Konrad Herbst, and Josh Goldford for their guidance and insight. Additionally, we would like to thank Corinne Vietorisz for assistance with soil community sampling. M.R.S. was supported by a synthetic biology NIH-funded predoctoral training fellowship (T32GM130546), a bioinformatics NIH-funded predoctoral training fellowship (T32GM100842), and the Biological Design Center Multicellular Design Program. J.M.B. was supported by DOE BER award DE-SC0020403. D.S. was supported by grants from the National Science Foundation (NSFOCE-BSF 1635070 and the NSF Center for Chemical Currencies of a Microbial Planet) and the U.S. Department of Energy, Office of Science, Office of Biological & Environmental Research through the Microbial Community Analysis and Functional Evaluation in Soils Science Focus Area Program (m-CAFEs) under contract number DE-AC02-05CH11231 to Lawrence Berkeley National Laboratory.

