## Supplement for "Metabolic complexity drives divergence in microbial communities"

**Supplementary material for "Metabolic complexity drives divergence in microbial communities"**

### Supplementary figures

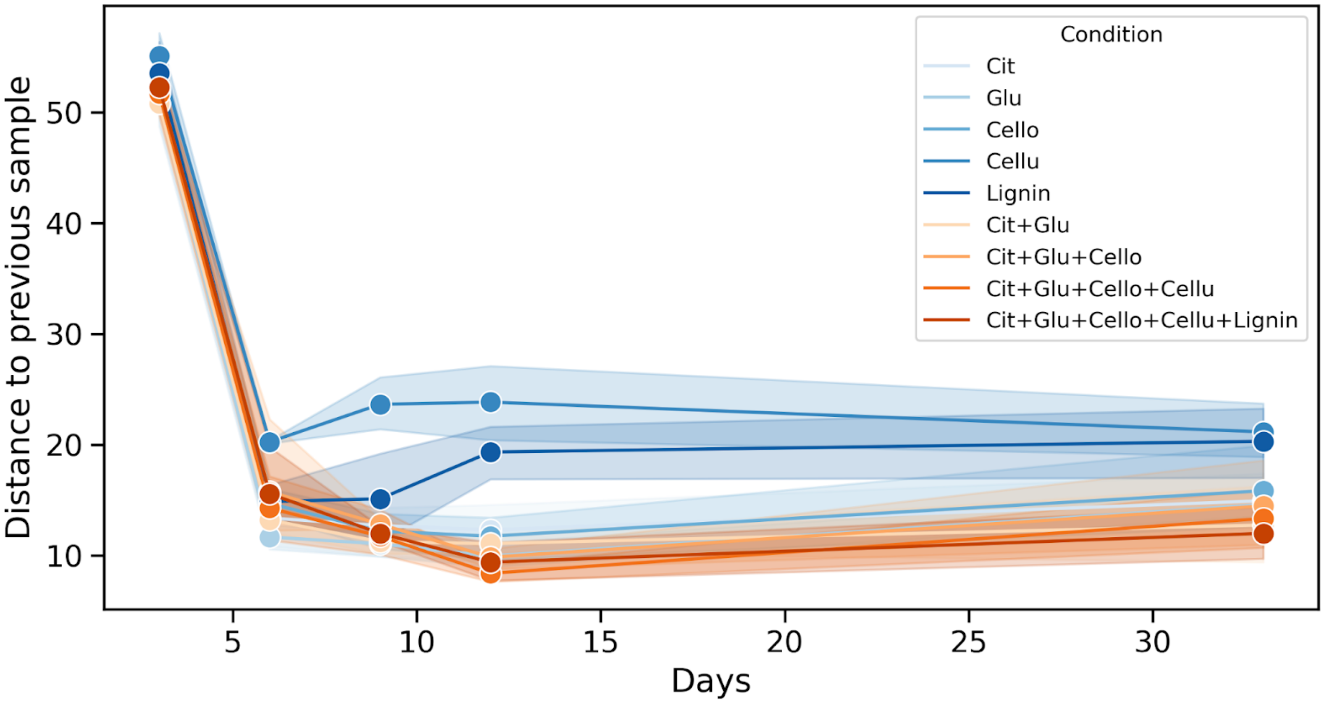


**Supplementary Figure 1 | Microbial communities stabilize.** Each point is the distance to a community's previous timepoint grouped by condition. On day 3, communities are substantially different from their initial state and then continue to change in composition, but by approximately the same amount.

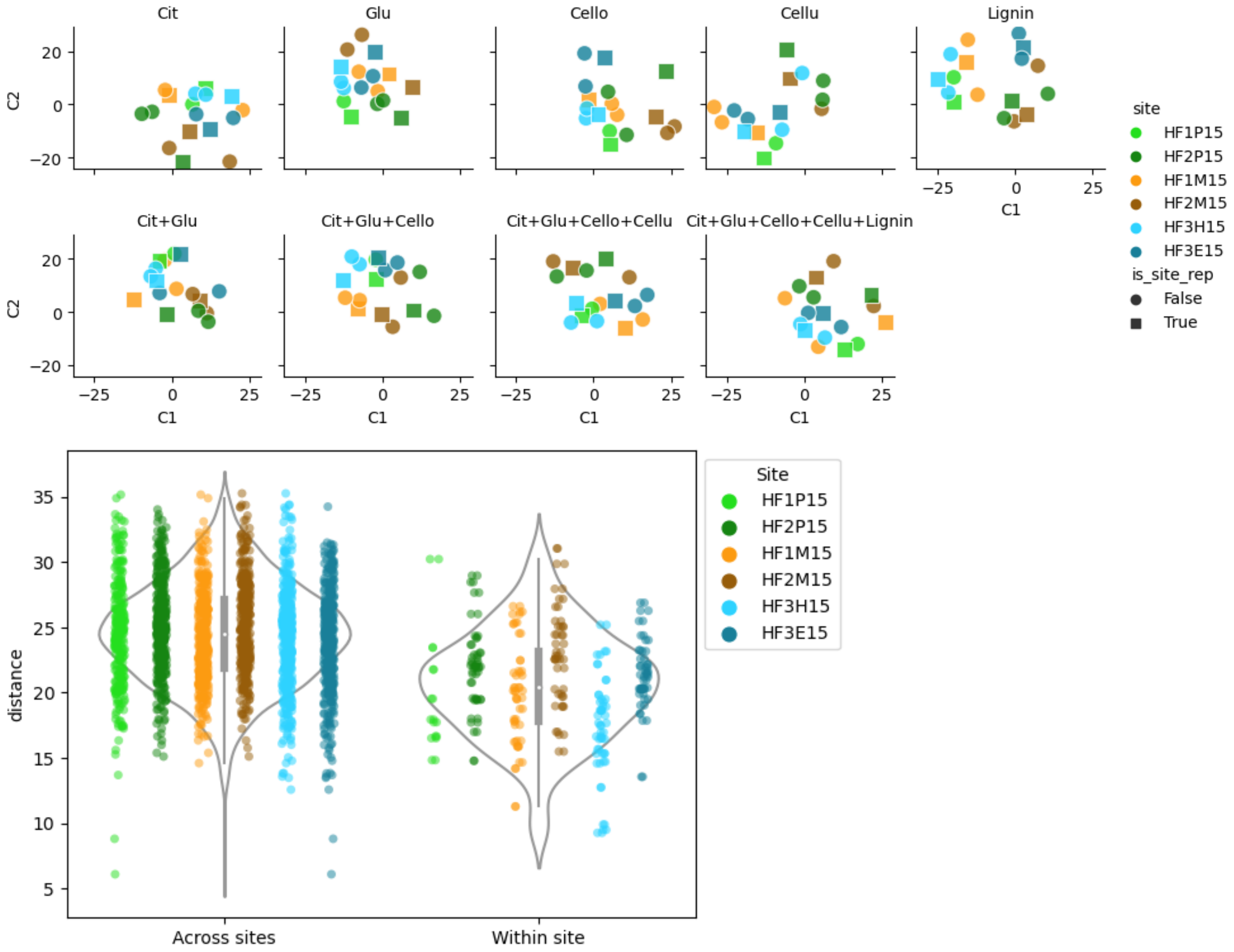


**Supplementary Figure 2 | Microcosm replicates cluster.**
The final state of three replicates for each community projected separately for each condition. Communities are colored by their source and squares represent the replicate used in the main text. Independent of condition, communities from the same replicate are more similar to each other than communities from separate replicates.

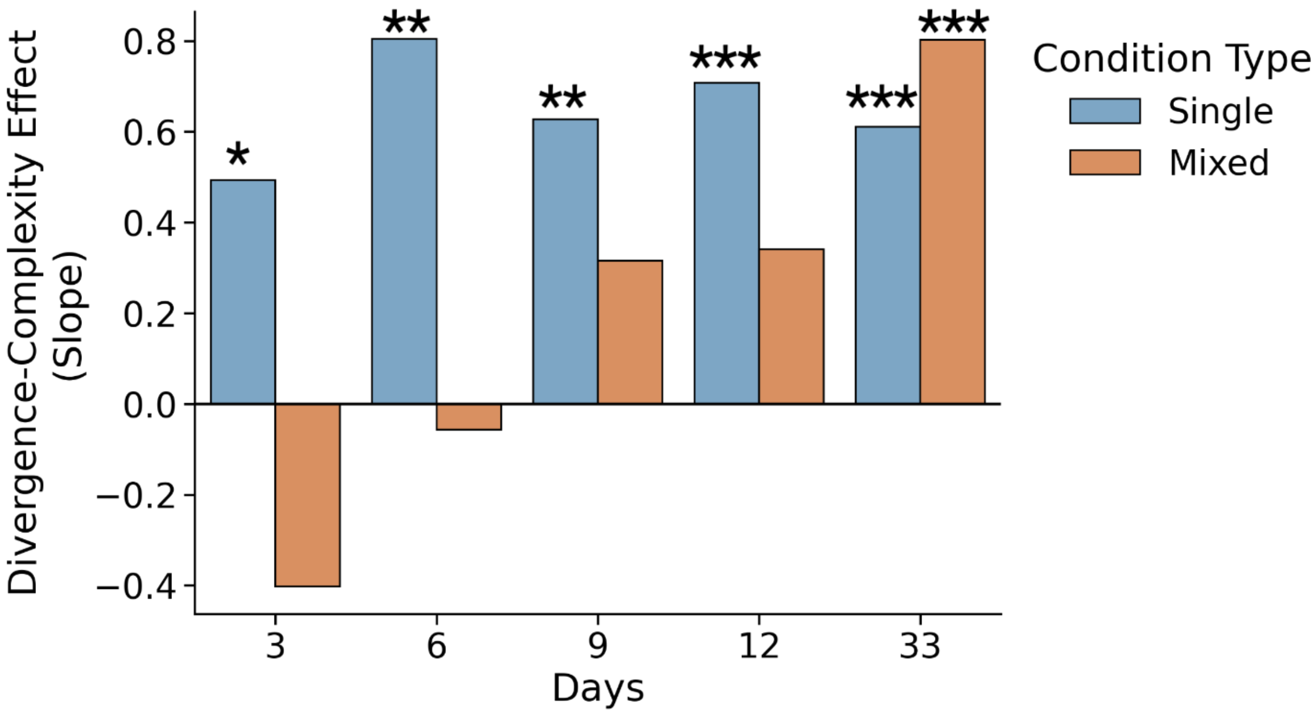


**Supplementary Figure 3 |** Same as Fig 2m but at Family level instead of ASV. *=p<.05, **=p<.01, ***=p<.001.

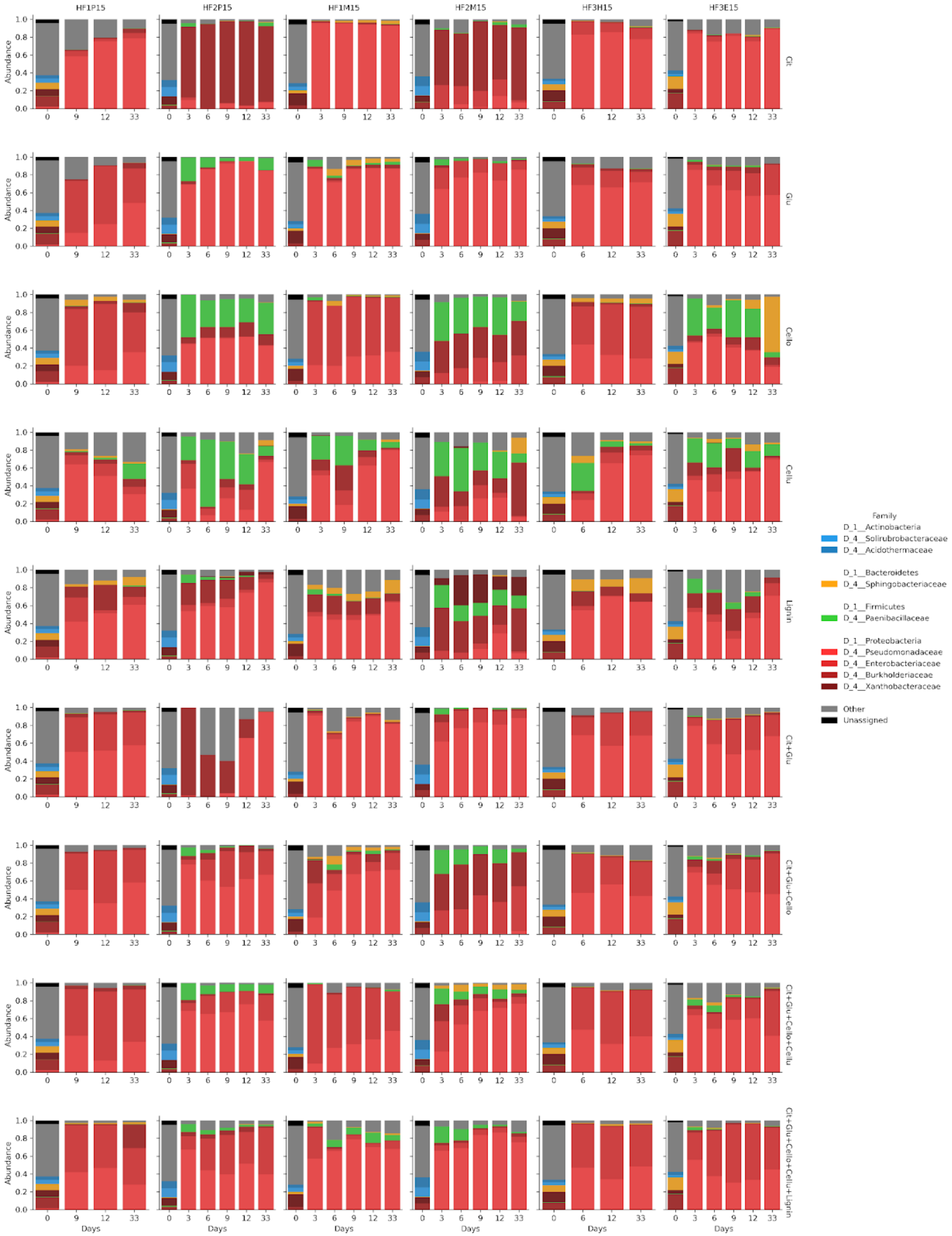


**Supplementary Figure 4 | Family composition over time.** Colors grouped by Phylum.

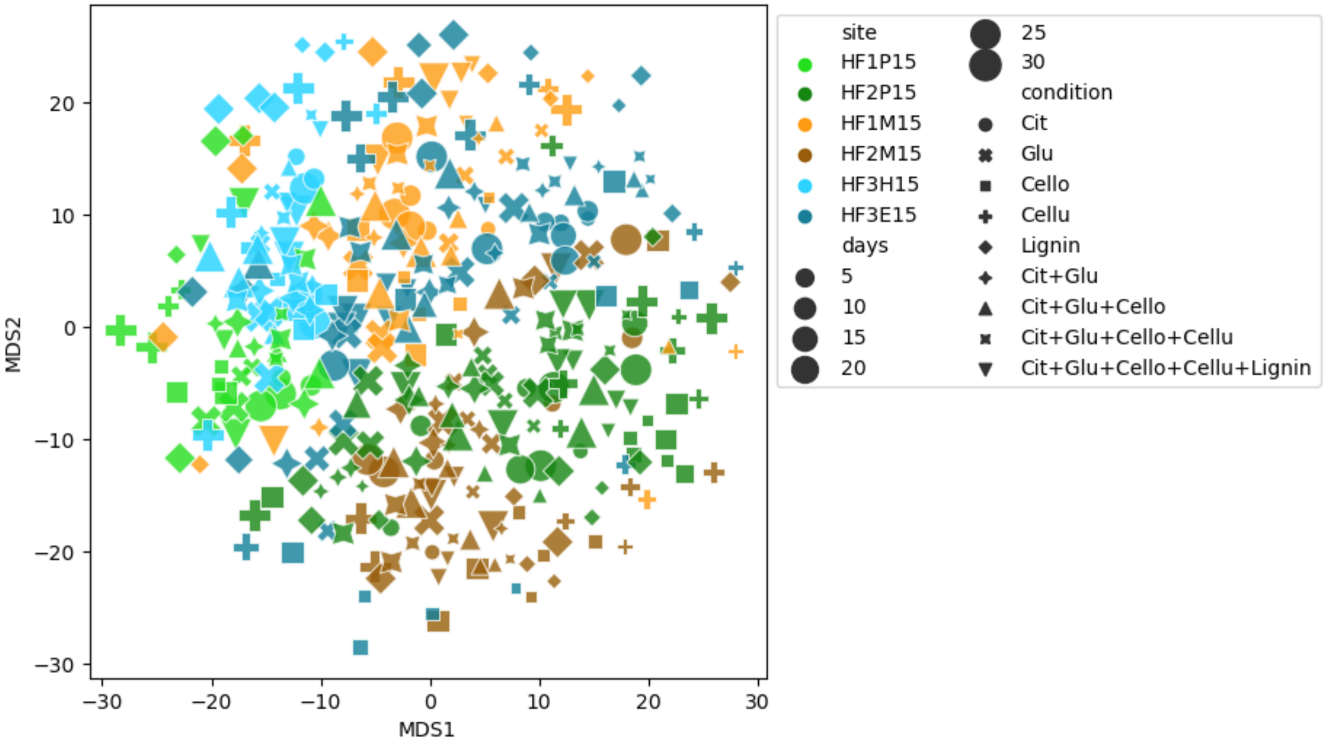


| Formula | $R^2$ |
|---|---|
| Distance ~ site * condition + days | 0.54 |
| Distance ~ site + days | 0.23 |
| Distance ~ condition + days | 0.14 |

**Supplementary Figure 5 | Overall, communities cluster more strongly by origin than by condition.** This is actually a pretty major result - it seems like most people think that environment is the strongest driver of community assembly, which these results challenge. Figure 2 is already busy, but this is related to this conclusion. Could just do the table in the main text and then have the figure as a supplement.

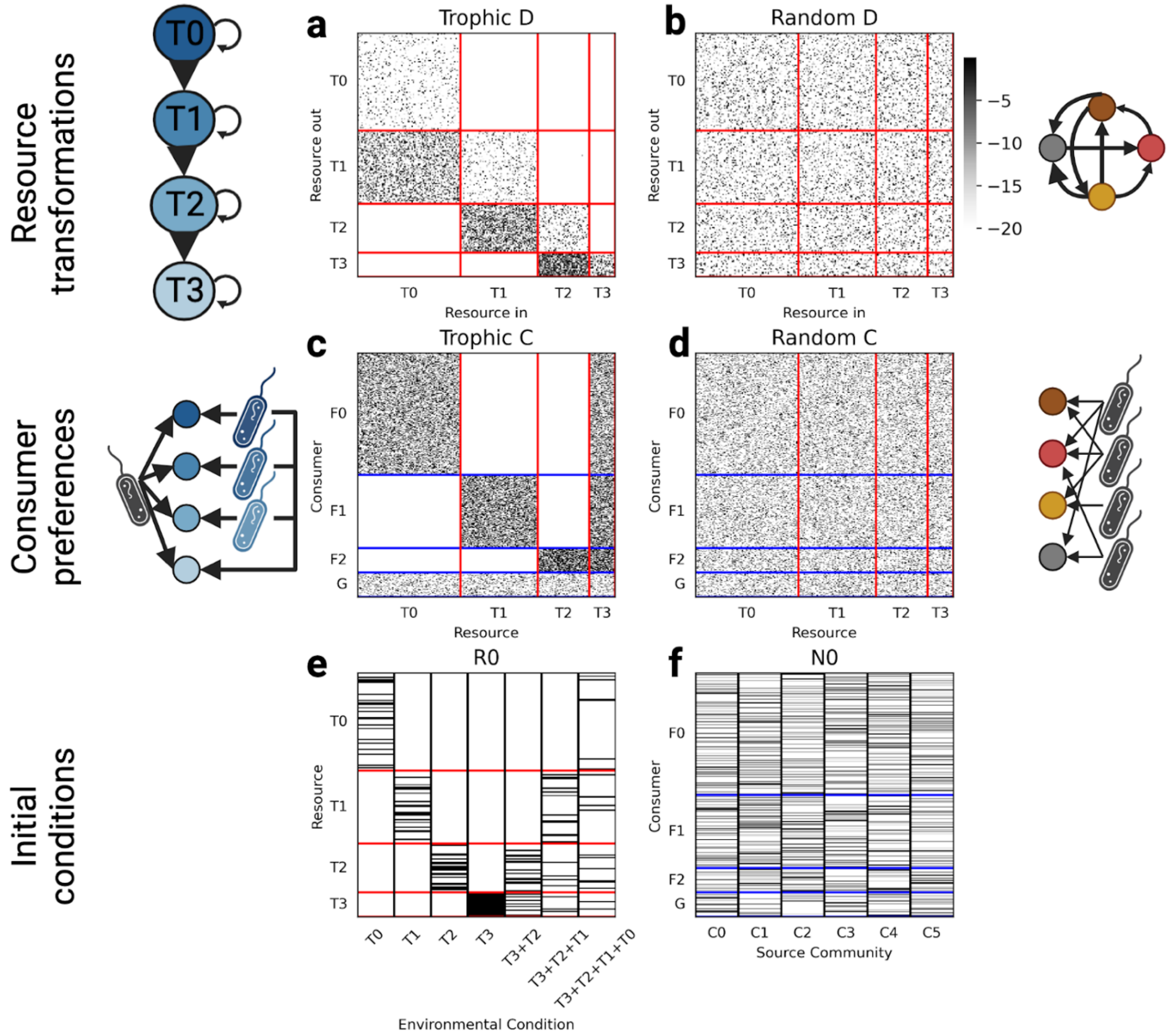


**Supplementary Figure 6 | Consumer resource parameterizations**. *D* Matrices for trophic (**a**) and random (**b**) resource transformations. Each column defines which resources are generated from a given input resource. **a,** Trophic resource transformations were parameterized by defining resource types (T0, T1, T2, and T3; red lines) which mostly transform into resources of the subsequent type with some self-renewal. **b,** Random resource transformations were parameterized by defining any resource to transform into any other resource with uniform probability. *C* matrices for trophic (**c**) and random (**d**) consumer preferences. Each row defines which resources can be utilized by each consumer. **c,** Trophic consumer preferences were parameterized by defining consumer families (F0, F1, F2, and G; blue lines) which consume a total of 35 resources of their associated type (for example, F0 consumers utilize T0 resources) and a common resource, T3. G consumers (generalists) consume resources of any type. **D,** Random consumer preferences were parameterized by defining consumers to utilize any resource with uniform probability. **e,** Seven initial environmental conditions were defined with 20 resources sampled from (1) T0, (2) T1, (3) T2, (4) T3, (5) T3+T2, (6) T3+T2+T1, and (7) T3+T2+T1+T0. **f**, Six source communities were defined by sampling 200 consumers.

| Day | Condition | HF1P | HF2P | HF3H | HF3E | HF1M | HF2M | Total |
|---|---|---|---|---|---|---|---|---|
| 0 | | 1/1 | 1/1 | 1/1 | 1/1 | 1/1 | 1/1 | 6 |
| 3 | Citrate | **0/1** | 1/1 | **0/1** | 1/1 | 1/1 | 1/1 | 4 |
| | Glucose | **0/1** | 1/1 | **0/1** | 1/1 | 1/1 | 1/1 | 4 |
| | Cellobiose | **0/1** | 1/1 | **0/1** | 1/1 | 1/1 | 1/1 | 4 |
| | Celluose | **0/1** | 1/1 | **0/1** | 1/1 | 1/1 | 1/1 | 4 |
| | Lignin | **0/1** | 1/1 | **0/1** | 1/1 | 1/1 | 1/1 | 4 |
| | C+G | **0/1** | 1/1 | **0/1** | 1/1 | 1/1 | 1/1 | 4 |
| | C+G+C | **0/1** | 1/1 | **0/1** | 1/1 | 1/1 | 1/1 | 4 |
| | C+G+C+C | **0/1** | 1/1 | **0/1** | 1/1 | 1/1 | 1/1 | 4 |
| | C+G+C+C+L | **0/1** | 1/1 | **0/1** | 1/1 | 1/1 | 1/1 | 4 |
| 6 | Citrate | **0/1** | 1/1 | 1/1 | 1/1 | **0/1** | 1/1 | 4 |
| | Glucose | **0/1** | 1/1 | 1/1 | 1/1 | 1/1 | 1/1 | 5 |
| | Cellobiose | **0/1** | 1/1 | 1/1 | 1/1 | 1/1 | 1/1 | 5 |
| | Celluose | **0/1** | 1/1 | 1/1 | 1/1 | **0/1** | 1/1 | 4 |
| | Lignin | **0/1** | 1/1 | 1/1 | 1/1 | 1/1 | 1/1 | 5 |
| | C+G | **0/1** | 1/1 | 1/1 | 1/1 | 1/1 | 1/1 | 5 |
| | C+G+C | **0/1** | 1/1 | 1/1 | 1/1 | 1/1 | 1/1 | 5 |
| | C+G+C+C | **0/1** | 1/1 | 1/1 | 1/1 | 1/1 | 1/1 | 5 |
| | C+G+C+C+L | **0/1** | 1/1 | 1/1 | 1/1 | 1/1 | 1/1 | 5 |
| 9 | Citrate | 1/1 | 1/1 | **0/1** | 1/1 | 1/1 | 1/1 | 5 |
| | Glucose | 1/1 | 1/1 | **0/1** | 1/1 | 1/1 | 1/1 | 5 |
| | Cellobiose | 1/1 | 1/1 | **0/1** | 1/1 | 1/1 | 1/1 | 5 |
| | Celluose | 1/1 | 1/1 | **0/1** | 1/1 | 1/1 | 1/1 | 5 |
| | Lignin | 1/1 | 1/1 | **0/1** | 1/1 | 1/1 | 1/1 | 5 |
| | C+G | 1/1 | 1/1 | **0/1** | 1/1 | 1/1 | 1/1 | 5 |
| | C+G+C | 1/1 | 1/1 | **0/1** | 1/1 | 1/1 | 1/1 | 5 |
| | C+G+C+C | 1/1 | 1/1 | **0/1** | 1/1 | 1/1 | 1/1 | 5 |
| | C+G+C+C+L | 1/1 | 1/1 | **0/1** | 1/1 | 1/1 | 1/1 | 5 |
| 12 | Citrate | 1/1 | 1/1 | 1/1 | 1/1 | 1/1 | 1/1 | 6 |
| | Glucose | 1/1 | 1/1 | 1/1 | 1/1 | 1/1 | 1/1 | 6 |
| | Cellobiose | 1/1 | 1/1 | 1/1 | 1/1 | 1/1 | 1/1 | 6 |
| | Celluose | 1/1 | 1/1 | 1/1 | 1/1 | 1/1 | 1/1 | 6 |

| | | | | | | | | |
|---|---|---|---|---|---|---|---|---|
| | Lignin | 1/1 | 1/1 | 1/1 | 1/1 | 1/1 | 1/1 | 6 |
| | C+G | 1/1 | 1/1 | 1/1 | 1/1 | 1/1 | 1/1 | 6 |
| | C+G+C | 1/1 | 1/1 | 1/1 | 1/1 | 1/1 | 1/1 | 6 |
| | C+G+C+C | 1/1 | 1/1 | 1/1 | 1/1 | 1/1 | 1/1 | 6 |
| | C+G+C+C+L | 1/1 | 1/1 | 1/1 | 1/1 | 1/1 | 1/1 | 6 |
| | Citrate | **2/3** | 3/3 | 3/3 | 3/3 | 3/3 | 3/3 | 17 |
| | Glucose | **2/3** | 3/3 | 3/3 | 3/3 | 3/3 | 3/3 | 17 |
| | Cellobiose | **2/3** | 3/3 | 3/3 | 3/3 | 3/3 | 3/3 | 17 |
| | Celluose | **2/3** | 3/3 | 3/3 | 3/3 | 3/3 | **2/3** | 16 |
| 33 | Lignin | **2/3** | 3/3 | 3/3 | 3/3 | 3/3 | 3/3 | 17 |
| | C+G | **2/3** | 3/3 | 3/3 | 3/3 | 3/3 | 3/3 | 17 |
| | C+G+C | **2/3** | 3/3 | 3/3 | 3/3 | 3/3 | 3/3 | 17 |
| | C+G+C+C | **2/3** | 3/3 | 3/3 | 3/3 | 3/3 | 3/3 | 17 |
| | C+G+C+C+L | **2/3** | 3/3 | 3/3 | 3/3 | 3/3 | 3/3 | 17 |
| | Total | 37 | 64 | 46 | 64 | 62 | 63 | 336 |

**Supplementary Table 1 | Availability of each sample**. When available, one sample was sequenced from each microcosm for days 0, 3, 6, 9, and 12. Three samples were sequenced for day 33. Bolded text indicates unsequenced conditions.
